# Bacteriophage populations mirror those of bacterial pathogens at sites of infection

**DOI:** 10.1101/2023.05.17.541184

**Authors:** NL Haddock, LJ Barkal, PL Bollyky

## Abstract

Bacteriophages, viruses that parasitize bacteria, are known to be abundant at sites of bacterial colonization but the relationship between phages and bacteria at sites of infection is unclear. Bacteriophage are highly specific to their bacterial host species and so we hypothesize that phage populations would mirror those of bacterial pathogens within infected tissues. To test this, here we study publicly-available cell-free DNA generated using next generation sequencing of infected bodily fluids, including urine, joint fluid, peritoneal fluid, bronchoalveolar lavage fluid, cerebrospinal fluid, and abscess fluid as well as uninfected control samples. These were analyzed using a computational pipeline for identifying bacteriophage sequences in cfDNA. We find that bacteriophage sequences are present in both infected and uninfected bodily fluids and represent a variety of bacteriophage morphologies and bacterial hosts. Additionally, phages from *E. coli, Streptococcus*, and *S. aureus* are overrepresented both in terms of proportion and diversity in fluids infected with these same pathogens. These data indicate that phages reflect the relative abundance of their bacterial hosts at sites of infection. Bacteriophage sequences may help inform future investigative and diagnostic approaches that utilize cell-free DNA to study the microbiome within infected tissues.

**Importance:** Bacteriophages are an active area of investigation in microbiome research but most studies have focused on phage populations at sites of bacterial colonization. Little is known about bacteriophage ecology at sites of active infection. To address this gap in knowledge, we utilized a publicly available dataset to study bacteriophage populations in cell free DNA collected from sites of infection. We find that phages reflect the relative abundance of their bacterial hosts at sites of infection. These studies may lead to future investigative and diagnostic approaches that incorporate phages as well as bacterial cell free DNA.

## Main Text

Bacterial infections are a major public health issue, causing morbidity and mortality worldwide. Recent studies indicating that bacterial infections may cause 13.6% of all global deaths^1^. In recent decades, the rise of antimicrobial resistance in bacterial infections has become increasingly common^2^ and poses a growing threat to vulnerable populations. The study of bacterial ecology at these sites is critical for understanding infection outcomes and developing improved diagnostics and therapeutics. However, much remains unknown.

Bacteriophage (phage) are viruses which infect bacteria, and they are highly specific to their bacterial hosts^3^. Phage are present ubiquitously throughout the human body^4,5^ and both reflect and influence bacterial populations. For this reason, phage are an attractive therapeutic target^6,7^, but outside of the study of phage therapy endogenous phages have been largely disregarded. Phage have been shown to be associated with bacterial host characteristics such as antimicrobial resistance^8^ or infection chronicity^9^. Little is known about phage ecology both in the human body as a whole but what is known is mostly relevant to sites of bacterial colonization such as in the gut^5,10^. Aside from recent work describing phage populations in chronically infected tissues^11^, the relationship between phages and bacteria at sites of infection is mostly unclear.

Here, we have investigated phage populations in infected bodily fluids and compared these versus uninfected controls. To accomplish this, we utilize next generation sequencing (NGS) data of cell-free DNA (cfDNA) from a publicly available study. cfDNA, though being largely comprised of human sequences, reflects microbial – and bacteriophage – sequences^12^. We utilize here a publicly available dataset of 76 infected bodily fluid samples (Bioproject PRJNA558701) generated via Illumina sequencing^13^. These included samples from infected wounds, joints, urine, serum, and other sites as well as culture negative controls. Causative infectious agents were identified by culture or 16S rRNA sequencing, including *Escherichia coli, Streptococcus spp*., and *Staphylococcus spp*. This dataset provides an excellent setting to investigate the relationships between pathogen and bacteriophage at the site of infection.

To identify phages within the metagenomic sequencing data, we apply a previously described bacteriophage annotation pipeline (Fig. 1A)^14^. In brief, raw data was quality controlled and trimmed, human host reads were subtracted by mapping to the human reference genome, and a BLAST search utilizing the full NCBI Nucleotide database was used to assess non-human cfDNA proportions. This revealed that on average, 8.8% of identifiable non-human reads were from bacteriophage in infected fluids, indicating that at the average site of infection bacteriophage comprise a considerable proportion of free DNA (Fig. 1B). Of note, an average of 12.8% of non-human reads belong to bacteriophage in the uninfected surgical control fluid samples – indicating that bacteriophage are abundant in these fluids independent of infection status. A first-pass BLAST search^15,16^ was performed our previously described Curated Phage Database (CPD) with stringent removal of sequences with human genome homology^14^. To link phage with bacterial host(s), we utilize a Curated Phage Dictionary^14^, naming convention of bacteriophages with clearly identified host genera, and NCBI nucleotide source host fields for bacteriophage sequence entries. This dictionary includes taxonomic classifications for both the phage and for the bacterial host, if known. Subsequent annotations reflect a diverse pool of bacteriophage across many families (Fig. 1C).

**Figure 1.**
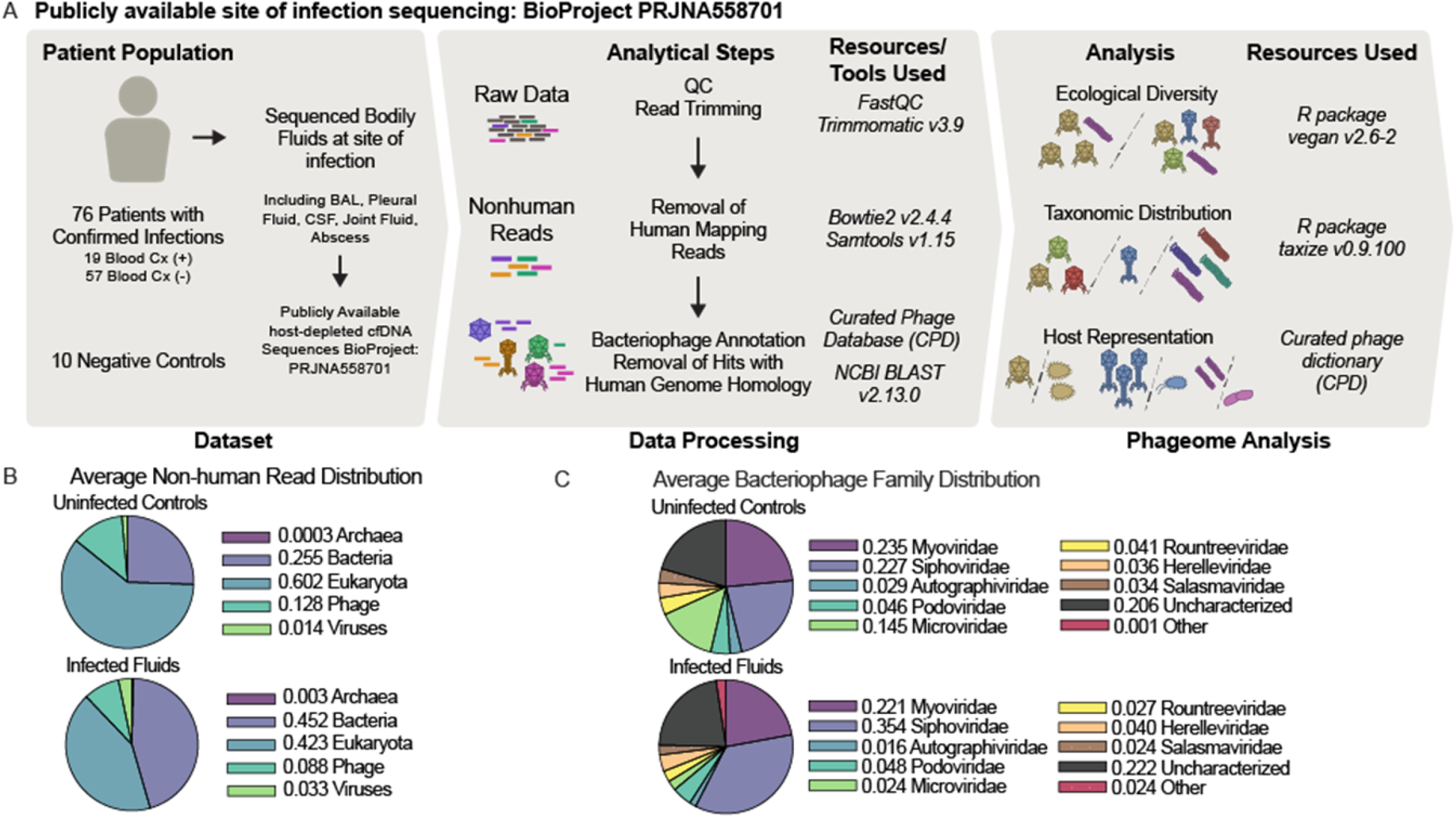
Infected and uninfected bodily fluids possess bacteriophage sequences A) Schematic of sample numbers, sample processing, and analytical approach. B) Distribution of non-human reads identity by Archeal, Eukaryotic, Bacterial, Mammalian Viral, and Bacteriophage categories. Uninfected controls: (Archaea mean 0.0003 SD 0.0005, Eukaryotic mean 0.602 SD 0214, Bacterial mean 0.255 SD 0.238, Mammalian Viral mean 0.014 SD 0.017, Bacteriophage mean 0.128 SD 0.102) Infected Fluids: (Archaea mean 0.003 SD 0.028, Eukaryotic mean 0.423 SD 0.330, Bacterial mean 0.452 SD 0.370. Mammalian Viral mean 0.033 SD 0.073. Bacteriophage mean 0.088 SD 0.111) C) Average bacteriophage distribution by bacteriophage family.

We find that when comparing overall composition of phageomes by bacterial host – there is notable enrichment for pathogen-specific phage by infection etiology, and that there is a non-zero phage background in the negative control samples which possessed *Escherichia, Klebsiella, Enterobacter*, and other phages (Fig. 2A). When analyzing these local phageomes with respect to infection etiology, we find that *E. coli* phage are increased in proportion compared to other infections (Fig. 2A,B). We find similar trends in *Streptococcus* and *Staphylococcus aureus* infections (Fig. 2A,C,D). Ecological diversity of phage, as calculated by the Shannon Diversity Index (SDI), a measure of entropy often applied in ecology to quantify species richness and evenness^17^, is disrupted on a per-infection basis (Fig. 2E-G), with higher levels of specific phage diversity in patients with infection by the corresponding host but not in controls or those with other infection etiology.

**Figure 2.**
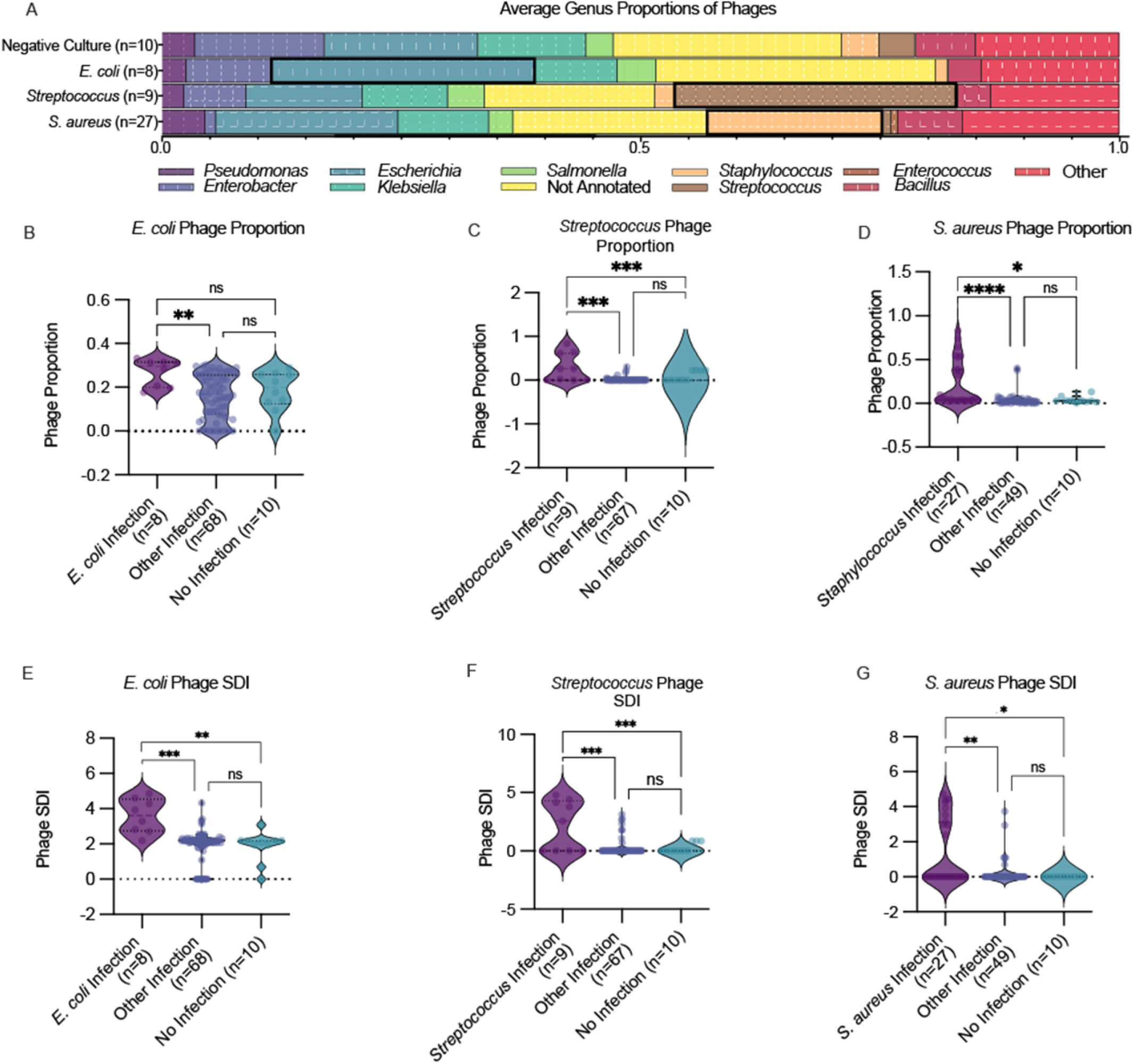
Local infections are associated with increased pathogen associated phage proportion and diversity. A) Average distribution of unique phages by bacterial host genus by infection category, with causative pathogen’s own bar outlined in black outline. (B-D) lnfection specific Phage Proportion calculated and shown as violin plots with median and quartiles shown by dashed lines for B) E. coli (Kruskal-Wallis P = 0.0083, mean E. coli Infection = 0.2666, mean Other Infection = 0.1575, mean No Infection =0.1842. Dunn’s Multiple comparisons: E. coli Infection vs other infection P = 0.0062, E. coli Infection vs No Infection P = 0.1456, Other Infection vs No Infection P > 0.9999), C) Streptococcus (Kruskal-Wallis P = 5.44E-5, mean Streptococcus Infection = 0.2939, mean Other Infection = 0.0145, mean No Infection = 0.00. Dunn’s Multiple comparisons: Streptococcus Infection vs other infection P = 0.0001, Streptococcus Infection vs No Infection P = 0.0002, Other Infection vs No Infection P = 0.8642), D) S. aureus (Kruskal-Wallis P = 3.14E-5, mean S. aureus Infection = 0.2215, mean Other Infection= 0.0418, mean No Infection= 0.0411. Dunn’s Multiple comparisons: S. aureus Infection vs other infection P = 1.98E-5, S. aureus Infection vs No Infection P = 0.0400, Other Infection vs No Infection P > 0.9999), E-G)Infection specific Shannon DI calculated and shown as violin plots with median and quartiles shown by dashed lines for E) E. coli (Kruskal-Wallis P = 0.0003, mean E.coli Infection Phage SDI = 3.591, mean Other Infection Phage SDI= 1.884, mean No Infection Phage SDI =1.854. Dunn’s Multiple comparisons: E. coli Infection vs other infection P = 0.0002, E. coli Infection vs No Infection P = 0.0018, Other Infection vs No Infection P > 0.9999), F) Streptococcus (Kruskal-Wallis P = 0.0003, mean Streptococcus Infection Phage SDI = 2.187, mean Other Infection Phage SDI = 0.2422, mean No Infection Phage SDI 0.00. Dum’s Multiple comparisons: Streptococcus Infection vs other infection P = 0.0002, Streptococcus Infection vs No Infection P = 0.0003, Other Infection vs No Infection P = 0.3549), G) S. aureus (Kruskal-Wallis P = 0.0036,mean S. aureus Infection Phage SDI = 1.362, mean Other Infection Phage SDI = 0.2178, mean No Infection Phage SDI 0.00. Dum’s Multiple comparisons: S. aureus Infection vs other infection P = 0.0093, S. aureus Infection vs No Infection P = 0.0223, Other Infection vs No Infection P > 0.9999)

Taken together, these findings demonstrate that bacteriophage populations reflect bacterial pathogens in infected bodily fluids.

However, this work possesses several limitations. It is unclear whether the sequences characterized here are from active bacteriophage particles, free phage DNA, or prophage DNA from bacterial hosts. Further studies are needed to understand the sources of these DNA as well as how they enter these bodily fluids. Another limitation is that this study utilizes publicly available cfDNA data taken from one patient cohort and may reflect geographically enriched features.

In summary, we find that bacteriophage sequences are present in both infected and uninfected bodily fluids and represent a variety of bacteriophage morphologies and bacterial hosts. Additionally, we demonstrate that infection etiology is reflected in infected bodily fluids through pathogen-associated phage proportion and diversity. Bacteriophage sequences may help inform future investigative and diagnostic approaches that utilize cell-free DNA to study the microbiome within infected tissues.

